# α-tubulin regulation by 5’ introns in *S. cerevisiae*

**DOI:** 10.1101/2023.06.22.546163

**Authors:** Linnea C. Wethekam, Jeffrey K. Moore

## Abstract

Across eukaryotic genomes, multiple α- and β-tubulin genes require regulation to ensure sufficient production of tubulin heterodimers. Features within these gene families that regulate expression remain underexplored. Here we investigate the role of the 5’ intron in regulating α-tubulin expression in *S. cerevisiae*. We find that the intron in the α-tubulin, *TUB1*, promotes α-tubulin expression and cell fitness during microtubule stress. The role of the *TUB1* intron depends on proximity to the *TUB1* promoter and sequence features that are distinct from the intron in the alternative α-tubulin isotype, *TUB3*. These results lead us to perform a screen to identify genes that act with the *TUB1* intron. We identified several genes involved in chromatin remodeling, α/β-tubulin heterodimer assembly, and the spindle assembly checkpoint. We propose a model where the *TUB1* intron promotes expression from the chromosomal locus, and that this may represent a conserved mechanism for tubulin regulation under conditions that require high levels of tubulin production.

**Article Summary:** α and β-tubulin proteins are encoded by families of genes that must be coordinately regulated to supply the αβ heterodimers that form microtubules. This study by Wethekam and Moore identifies a role for the early intron in the budding yeast α-tubulin, TUB1, in promoting gene function. A genetic screen reveals new tubulin regulators that act through the TUB1 intron. The results establish new layers of α-tubulin regulation that may be conserved across eukaryotes.

## Introduction

Tubulin, the fundamental subunit of the microtubule cytoskeleton, is a heterodimeric protein consisting of α and β monomers. In many eukaryotes, the α and β monomers are encoded by a family of genes at disparate genomic loci. For example, in humans there are 9 well-annotated genes that encode α-tubulin and 8 β-tubulin genes, located across 5 and 7 chromosomes, respectively (Findeisen et al., 2014; Leandro-García et al., 2010; Park et al., 2021). Some of the tubulin genes are clustered together on chromosomes (for example: *TUBA1A, TUBA1B, TUBA1C*); however, many are the only tubulin gene encoded on its chromosome. It is unclear how the structure of the tubulin genes impacts tubulin expression.

Our previous work showed that cells must maintain excess α-tubulin for efficient mitotic timing and to prevent the accumulation of monomeric β-tubulin (Wethekam & Moore, 2023). Conserved features within coding sequences of α-tubulin genes may provide a point of regulation. One conserved feature of α-tubulin genes is an intron located near the 5’ end of the coding sequence (Figure 1A). Within the human α-tubulins, the early intron is positioned immediately 3’ of the start codon, while in both budding yeast α-tubulin genes, known as *TUB1* and *TUB3*, the intron is positioned 23 base pairs 3’ of the start codon (ORF; Figure 1A; Findeisen et al., 2014). The role of the early intron in α-tubulin gene function is unexplored.

**Figure 1.**
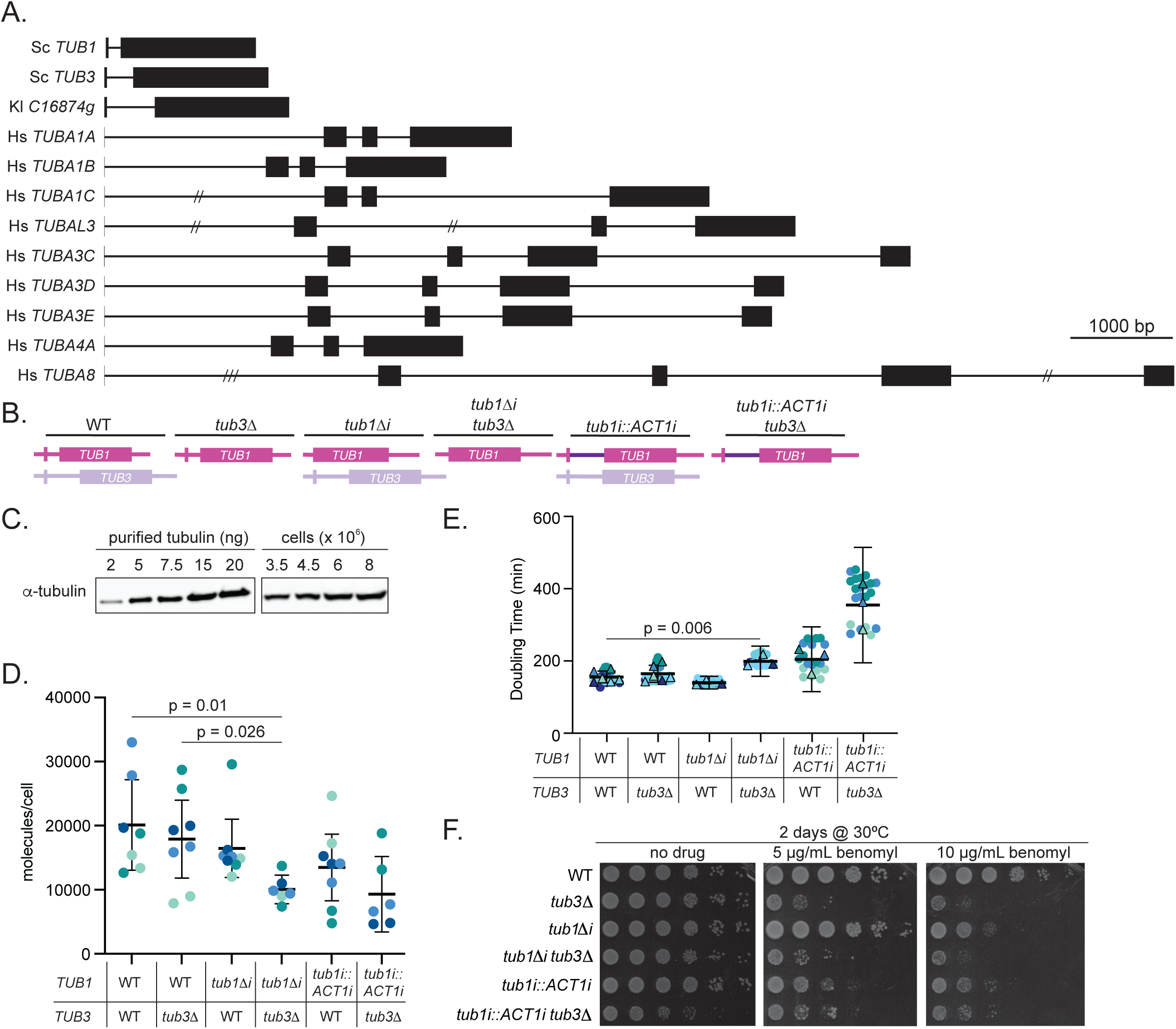
*TUB1* intron promotes α-tubulin expression. A. Diagram of α-tubulin genes from *S. cerevisiae*, *K. lactis*, and *H. sapiens*. All genes are drawn to scale with 20 bp/pixel. For five introns the scale was adjusted, // = 40 bp per pixel, /// = 80 bp per pixel. B. Diagram of genotypes used in this figure. C. Example western blot used to determine the number of α-tubulin molecules per cell. Left shows the purified tubulin standards used to build the standard curve. Right shows the estimated number of cells loaded onto the gel used to identify the number of molecules of α-tubulin per cell D. Quantification of α-tubulin molecules per cell across each genotype. Band intensities were converted to protein mass (ng) and then converted to molecules per cell. Data represent at least three independent experiments with two biological replicates per experiment. Each dot represents one mean molecules per cell measurement for at least three dilutions per biological replicate. Dots are colored by experiment E. Quantification of doubling times for the indicated mutants normalized to the mean of the technical replicates for wild type. For each genotype at least three technical replicates were used in at least three independent experiments. Each circle represents a single technical replicate, with triangles representing the mean of all technical replicates. Shapes are colored based on experiment. Bars represent mean ± 95% CI. P-values are from a t test comparing wild type and mutant, specific p-values are listed in the text. F. 10-fold dilution series of listed strains spotted onto rich media or rich media supplemented with 5 or 10 µg/mL benomyl. Plates were grown at 30°C for two days before imaging.

The conservation of the early intron in budding yeast α-tubulins is particularly notable. While introns are common within metazoan genomes, only 5% of the *S. cerevisiae* genes contain introns (Neuvéglise et al., 2011; Stajich et al., 2007). Of the intron containing genes in *S. cerevisiae*, a majority encode ribosomal proteins and removing the intron disrupts ribosome function and ribosomal protein expression, particularly under stress (Parenteau et al., 2011). Another example of a gene with a 5’ intron important for regulating protein production in *S. cerevisiae* is *ACT1*, the gene encoding g-actin. *ACT1* is the only gene that encodes g-actin in *S.* cerevisiae and must be expressed at high levels to maintain the dynamic actin cytoskeleton (Blank et al., 2020; Wertman et al., 1992). The *ACT1* intron has been shown to promote the expression of *ACT1* and is especially important when cells are stressed with the actin-depolymerizing drug, latrunculin A (Agarwal & Ansari, 2016; Juneau et al., 2006). Removal of the *ACT1* intron also reduces the amount of *ACT1* mRNA (Agarwal & Ansari, 2016; Juneau et al., 2006). Conversely, exogenously inserting the *ACT1* intron into another gene can increase the expression of that gene and this depends on the location of the intron within the coding sequence, with the enhancing capability of the intron decreasing the further it is placed 3’ of the the start codon (Agarwal & Ansari, 2016; Dwyer et al., 2021). This suggests that the α-tubulin early introns may also promote protein expression through a related mechanism.

In this study, we sought to understand the role of the α-tubulin introns in regulating α-tubulin protein production. We used budding yeast which contains two genes for α-tubulin, *TUB1* and *TUB3*, and both genes contain introns. We find that the intron in *TUB1* is important for promoting the expression of α-tubulin and resistance to microtubule stress. Furthermore, the *TUB1* intron exhibits a stronger effect than the *TUB3* intron, and partially rescues the stress sensitivity of cells expressing *TUB3* only. A genetic screen for potential regulators of *TUB1* that act through the intron identified genes encoding RNA-regulating proteins and chromatin modifiers, representing novel regulators of α-tubulin expression. Our screen also identified several genes known to be involved in heterodimer biogenesis and turnover, suggesting that the *TUB1* intron is important for the maintaining balance between heterodimer production and destruction.

## Results

### *TUB1* intron promotes α-tubulin expression

5’ introns are a common feature among α-tubulin genes. Both the *TUB1* and *TUB3* genes in *S. cerevisiae* contain single introns that begin 23 base pairs 3’ of the start codon (Figure 1A). A similar intron is found in the single α-tubulin of the budding yeast *K. lactis*, suggesting that the intron pre-dates the whole genome duplication (Figure 1A). Furthermore, all human α-tubulin genes exhibit an intron immediately 3’ of the start codon (Figure 1A). The prevalence of 5’ introns raises the hypothesis that these may represent an important and conserved feature for promoting α-tubulin gene function.

We used three experiments to determine if the *TUB1* intron is important for promoting α-tubulin expression in *S. cerevisiae*. First, we tested if the *TUB1* intron impacts α-tubulin at the protein level by performing quantitative western blots. To do this we used purified yeast tubulin to build standard curves of α-tubulin. Band intensities of cell lysates from log-phase cells grown in rich media were compared to the standard curve and converted from nanograms of protein to molecules per cell (see Material and Methods; Wethekam & Moore, 2023). In cells where we removed the *TUB1* intron, the level of α-tubulin across experiments and biological replicates was more consistent and tended to be less than what we observed in wild-type controls (Figure 1B-D). However, this apparent difference was not significant (p = 0.30). To isolate the output from the intron-less *tub1*Δi gene we knocked out the other α-tubulin isotype, *TUB3*. These *tub1*Δi *tub3*Δ cells exhibit approximately 50% less α-tubulin than wild-type controls or *TUB1 tub3*Δ cells (p = 0.01, p = 0.026; Figure 1B-D). These results suggest that the *TUB1* intron promotes α-tubulin protein production, and that this role is most apparent when *TUB1* is the only source of α-tubulin.

We next tested whether the *TUB1* intron is important for cell fitness by comparing doubling time in cells with or without the *TUB1* intron. Cells were grown to saturation, diluted 500-fold into rich media, and the OD_600_ was recorded every five minutes for 20 hours. We used these measurements to estimate the doubling time during the exponential phase of growth. In cells where we removed the *TUB1* intron, there is no distinguishable difference in doubling time between wild-type and *tub1Δi* cells (p = 0.15, Figure 1B, E). We also find no distinguishable difference in *tub3Δ* compared to wild type (p = 0.44, Figure 1B, E). In contrast we see a significant increase in the doubling time in *tub1Δi tub3Δ* cells, where the only source of α-tubulin is *tub1Δi* (p = 0.006, Figure 1B, E). Together these results indicate that the *TUB1* intron promotes fitness.

As a final test, we determined if the *TUB1* intron was important for tolerance of microtubule stress. Benomyl is a fungicide that creates microtubule stress by inhibiting tubulin polymerization (Gupta et al., 2004; Rathinasamy & Panda, 2006). We predicted that if removing the *TUB1* intron weakens α-tubulin production, then these cells should exhibit increased sensitivity to benomyl. Consistent with this prediction, *tub3Δ* cells that lack the minor α-tubulin isotype proliferate slower than wild-type control cells at low concentrations of benomyl (5 µg/mL; Figure 1B, F). *tub1Δi* cells are also hypersensitive to benomyl, but to a lesser degree than *tub3*Δ cells (Figure 1B, F). In *tub1Δi tub3Δ* double mutant cells exhibit a level of sensitivity that is similar to *tub3*Δ single mutant cells (Figure 1B, F). These results suggest that the *TUB1* intron is important for tolerance of microtubule stress.

### *TUB1* intron is not rescued by the *ACT1* intron

If the intron promotes *TUB1* function, we predicted that this function might be rescued by substituting an alternative intron that is known to promote expression of other genes. We tested this prediction by replacing the *TUB1* intron with the *ACT1* intron sequence at the endogenous *TUB1* locus, creating *tub1i∷ACT1i* (Figure 1B), and used the three experiments described above. We find that *tub1i∷act1i* cells hold approximately 33% less α-tubulin than wild type, and *tub1i∷ACT1i tub3Δ* double mutants hold 54% less α-tubulin than wild type (p = 0.08 and 0.02 respectively; Figure 1B-D). In the doubling time assay, *tub1i∷ACT1i* cells grow 20% slower than wild-type controls or *tub1Δi* cells (157.3 min versus 206.4 min; p = 0.02; Figure 1B, E). *tub1i::ACT1i tub3*Δ proliferate >100% slower than *tub3Δ* cells (356.5 min compared to 157.3 min; p = 0.0001; Figure 1B, E). In the benomyl assay, *tub1i∷ACT1i* cells exhibit greater sensitivity to benomyl than the wild-type controls or *tub1Δi* cells (Figure 1B, D). This is sensitivity is exacerbated in *tub1i∷ACT1i tub3Δ* double mutant cells (Figure 1B, D). Together these results indicate that the *ACT1* intron cannot rescue the *TUB1* intron, and suggest that the sequence of the *TUB1* intron contributes a distinct function.

### Comparing the introns of *TUB1* and *TUB3* isotypes

The introns in both *TUB1* and *TUB3* begin at the same position in the ORF but differ in sequence and total length (Figure 1A, 2A). The *TUB1* intron is 118bp long, and the *TUB3* intron is 300bp long (Figure 2A, S1). To test whether the *TUB1* and *TUB3* introns are functionally equivalent, we generated three chimeric alleles at the *TUB1* locus: 1) *tub1i::TUB3i*, which has the coding sequence of *TUB1* and the intron of *TUB3*; 2) *tub1Δ::TUB3*, which has the coding sequence and intron of *TUB3*; 3) *tub1Δ::TUB3^TUB1i^* which has the coding sequence of *TUB3* and the intron of *TUB1* (Figure 2B). Replacing the *TUB1* intron with that of *TUB3* showed a moderate level of benomyl sensitivity that matches the sensitivity of *tub1Δi* cells (Figure 2C). Consistent with prior work, *tub1∷ΔTUB3* cells are hypersensitive to benomyl (Figure 2B, C; Nsamba et al., 2021), but that benomyl sensitivity is partially rescued in *tub1Δ::TUB3^TUB1i^*cells (Figure 2B, C). This suggests that the *TUB1* intron provides a higher level of function than the *TUB3* intron, even when combined with the coding sequence of *TUB3.* To confirm the partial rescue in a separate assay, we measured doubling time in the presence of a different microtubule-destabilizing drug, nocodazole. Whereas the chimeric alleles show no effect in DMSO controls, in 5µM nocodazole *tub1Δ∷TUB3* cells show a strong increase in doubling time (248.7 min) and *tub1Δ∷TUB3^TUB1i^*cells show an intermediate effect (210.5 min), compared to wild-type controls (163.5 min; Figure 2D). Together these results suggest that the *TUB1* and *TUB3* introns are not functionally equivalent, and that the *TUB1* intron may promote a higher level of α-tubulin function.

**Figure 2.**
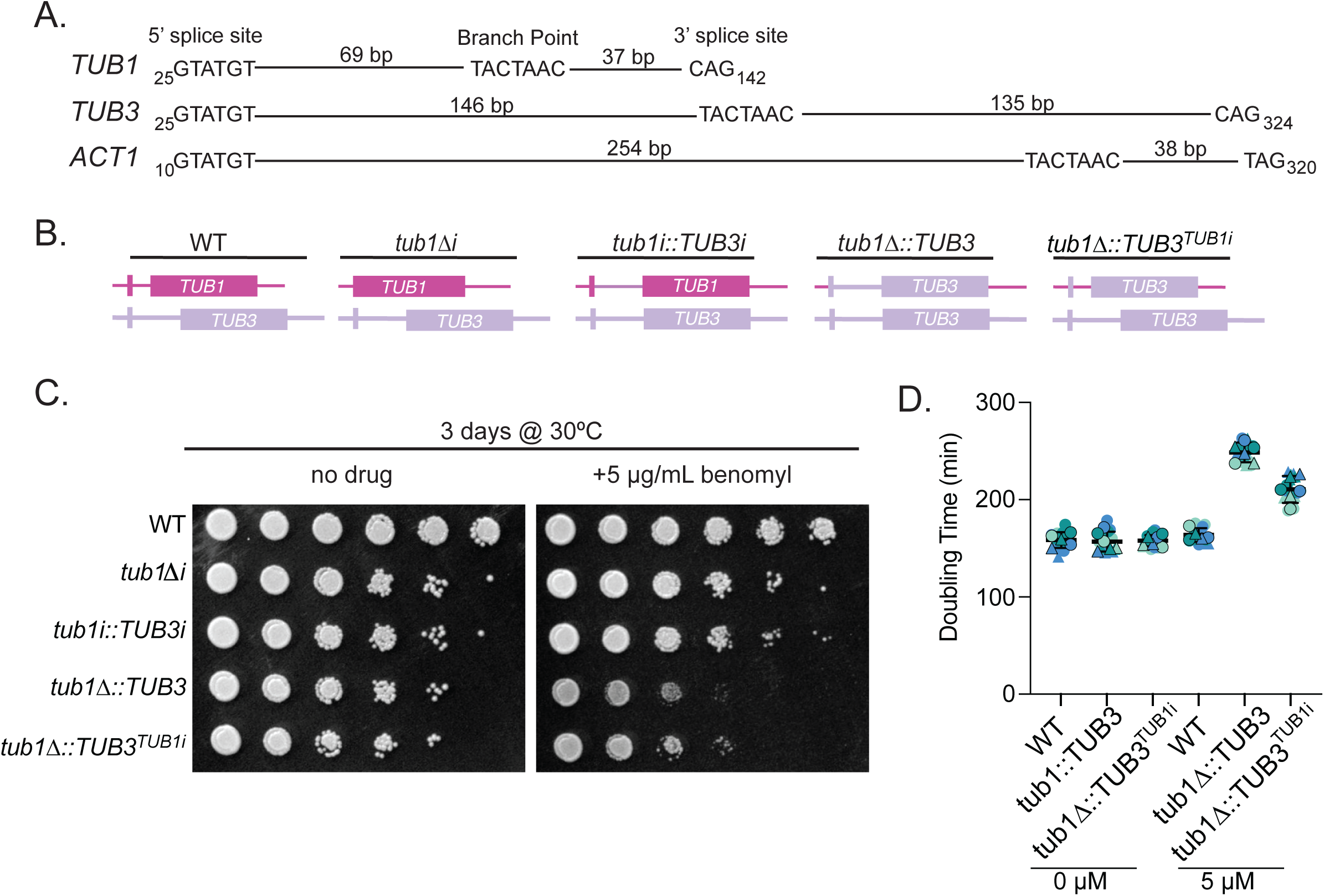
*TUB1* intron protects cells from microtubule stress. A. Diagram of both α-tubulin introns and the *ACT1* intron. The last base pair in exon 1 is listed before the 5’ splice site sequence. The length from the end of the 5’ splice site to beginning of the branch point sequence is listed and drawn to scale. Length from the end of the branch point sequence to the start of the 3’ splice site is also drawn to scale. The base pair of the exon 2 is listed immediately following the 3’ splice site. Scale is 1 bp per pixel. B. Diagram of the genotypes used in the benomyl sensitivity assay. C. 10-fold dilution series of listed strains spotted onto rich media or rich media supplemented with 5 µg/mL benomyl. Plates were grown at 30°C for two days before imaging. D. Quantification of doubling time for each of the indicated strain under indicated concentration of nocodazole. For each genotype two biological replicates were used in at least two independent experiments. Three technical replicates were used in each independent experiment. Circles represent one biological replicate and triangles represent the other biological replicate. Circles or triangles with boarders represent the means of the technical replicates, all other circles or triangles represent technical replicates.

### Identifying genes that act through the *TUB1* intron

Our results above indicate that the *TUB1* intron promotes α-tubulin expression. To elucidate the underlying mechanism and identify extrinsic regulators that might act in the same pathway with the intron, we used a genetic interaction screen with the collection of ∼5000 non-essential gene deletion strains (see Materials and Methods). We first predicted that loss of a gene that promotes *TUB1* expression would create hypersensitivity to benomyl, similar to cells that lack the *TUB1* intron (Figure 1D). We compared the growth of the haploid deletion collection on rich media supplemented with 10 µg/mL benomyl and 2% DMSO to rich media with 2% DMSO alone, and used a previously published image analysis method to measure and score the growth of four technical replicates of each strain (Wagih et al., 2013; Wagih & Parts, 2014). This analysis identified 649 gene deletions that exhibit hypersensitivity to benomyl (Table S1). To narrow this list and identify genes that may act in a pathway with the *TUB1* intron to promote α-tubulin expression, we focused on genes that are known to exhibit a negative genetic interaction with a *tub3Δ*, since loss of *TUB3* exacerbates the *tub1Δi* allele in our experiments. We identified 150 genes listed as negative genetic interactors with *tub3Δ* in the Saccharomyces Genome Database; 42 of these genes are also benomyl sensitive in our assay (Table S2). Finally, we predicted that genes acting through the *TUB1* intron would exhibit a positive genetic interaction with *tub1Δi*. In other words, combining a mutant allele that ablates gene function with a *TUB1* mutant allele that lacks its intron would exhibit a level of benomyl sensitivity that is equivalent or better than either single mutant alone. To identify this set of genes we generated double mutants by synthetic genetic array (see Materials and Methods) and quantitatively compared the growth of double mutants on 10 µg/mL benomyl with 2% DMSO to that of single mutants under the same conditions. We identified 33 genes where combining the deletion allele with the *tub1Δi* allele exhibits a similar level of benomyl sensitivity or improves benomyl sensitivity compared to the deletion allele alone (Table S2).

We performed a GO Term analysis to determine if the 33 genes we identified are known to act in shared pathways or processes (see Materials and Methods). We identified significantly enriched Cellular Component GO Terms associated with the Swr1 complex (*ARP6, SWC3, SWR1, TAF15, VPS71, YAF9*), prefoldin complex (*GIM4, GIM5, PAC10, YKE2*), microtubules (*CIN1, MAD2, NIP100, PAC2, TMA19, TUB3*), the bub1-bub3 complex (*BUB1, BUB3*), and the mitotic checkpoint complex (*BUB3, MAD2*; Figure 3A). To determine if any of our identified genes encode proteins that work together in complexes, we used GeneMania to map physical interactions between these gene products (Figure 3B; Montojo et al., 2010). This analysis identified components of the Swr1/Ino80 complex that replaces dimers of H2A-H2B histones for Htz1-H2B dimers, the GimC/Prefoldin complex that folds nascent α- and β-tubulin monomers, the tubulin binding cofactors (TBCs) that assemble tubulin monomers into heterodimers, and the spindle assembly checkpoint complex that prevents anaphase onset in the presence of mitotic spindle errors (Hansen et al., 1999; Lewis et al., 1997; Mizuguchi et al., 2004; Rudner & Murray, 1996; Tian et al., 1999; Vainberg et al., 1998; Figure 3B). This analysis suggests that genes identified in our screen are likely involved in promoting α-tubulin expression through altering the chromatin landscape or impacting tubulin folding and heterodimer assembly.

**Figure 3.**
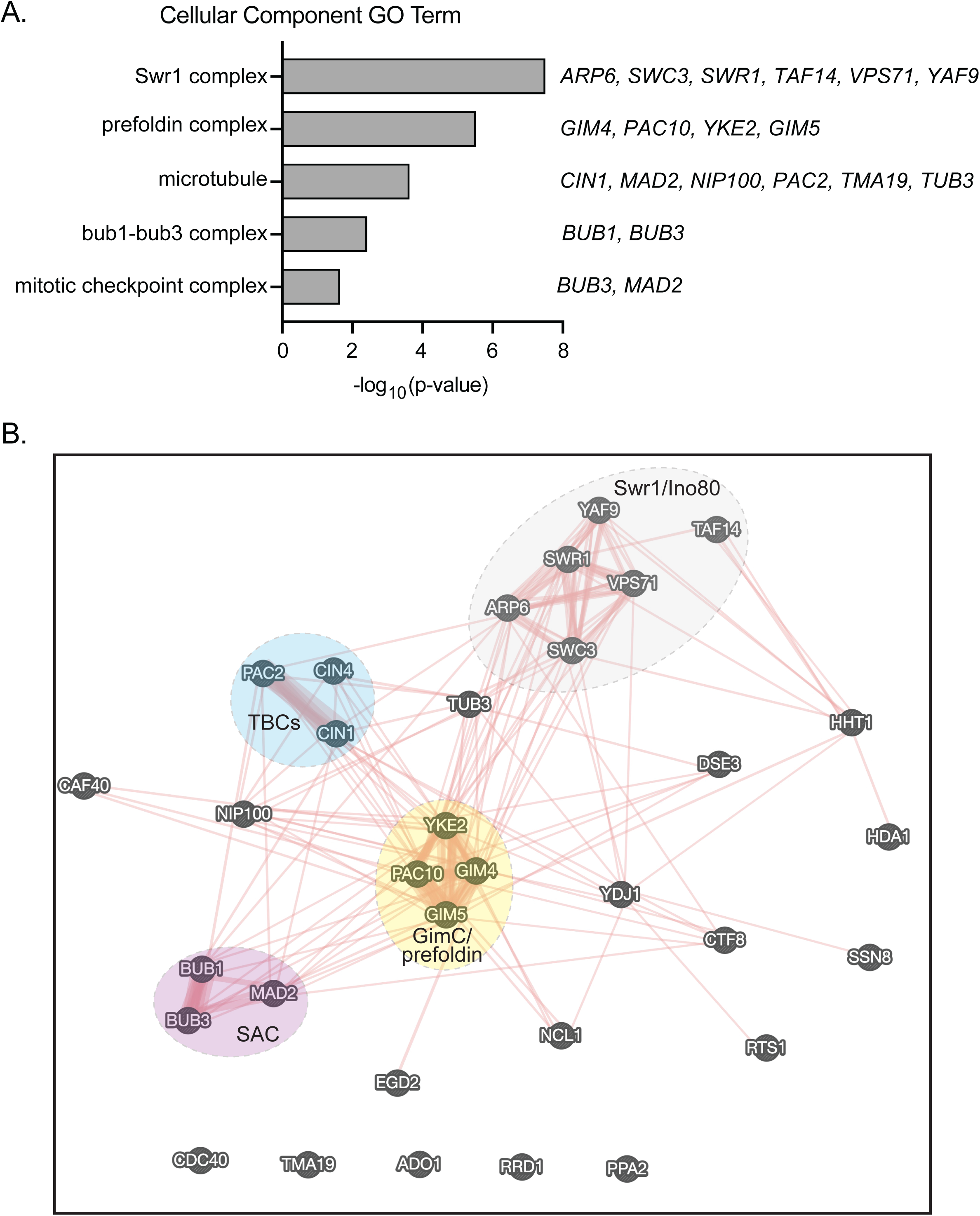
Genome-wide screen to identify genes that regulate *TUB1* through its intron. A. Plot of Cellular Component GO Terms with p < 0.05 and not redundant with other GO Terms. B. GeneMANIA network analysis of physical interactions between the 33 genes identified from our screening and filtering.

### Novel regulators of α-tubulin expression

We selected several genes from our screen for further investigation, based on their reported roles in RNA binding (*NCL1*) or chromatin regulation (*SWC3* and *VPS71*; Figure 3A). We also included *GIM5* since it has a well-established role in α-tubulin protein folding (Lacefield & Solomon, 2003; Vainberg et al., 1998). To confirm that loss of these genes diminishes α-tubulin expression, we tested whether adding low copy number plasmids expressing either *TUB1* or *TUB3* can suppress the benomyl sensitivity phenotype of the null mutants. Transformants were grown in media selective for the plasmid and then spotted to rich media or rich media containing benomyl. In control experiments, additional copies of *TUB1* or *TUB3* confer benomyl resistance to wild-type cells and rescue the benomyl sensitivity of *tub3*Δ mutants (Figure 4A, B). Additional copies of *TUB1* or *TUB3* also rescue the benomyl sensitivity of the *ncl1Δ*, *swc3Δ*, and *vps71Δ* mutants; but do not rescue the benomyl sensitivity of *gim5Δ* mutants (Figure 4A, B). These results demonstrate that increasing the copy number of α-tubulin genes can rescue the sensitivity of *ncl1Δ*, *swc3Δ*, and *vps71Δ* mutants to microtubule stress.

**Figure 4.**
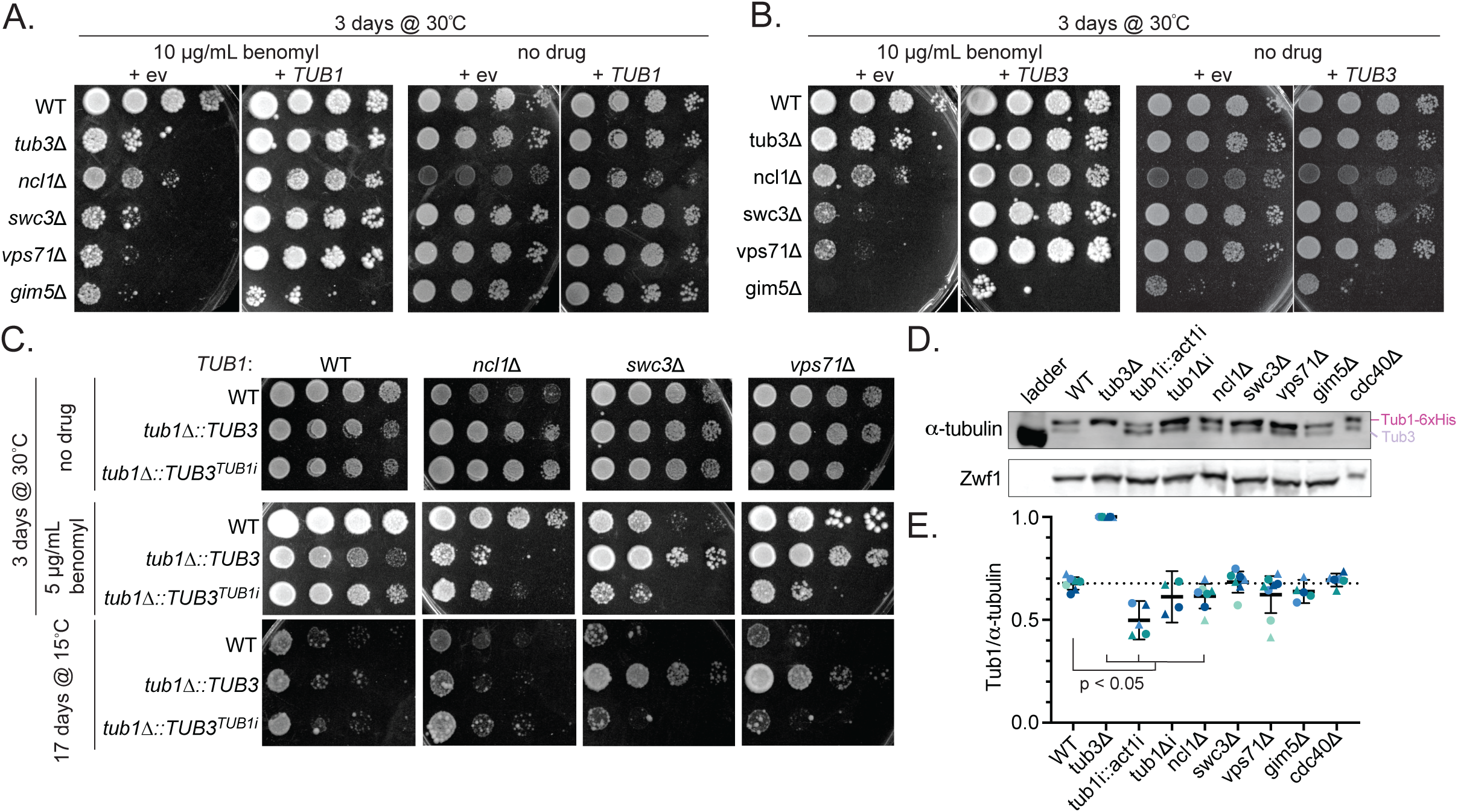
Screen hits identify novel regulators of α-tubulin expression. A. 10-fold dilution series of screen hits with an extra copy of *TUB1* or a control plasmid spotted onto rich media or rich media supplemented with 10 µg/ml benomyl. Plates were incubated at 30°C for 3 days before imaging. B. 10-fold dilution series of screen hits with an extra copy of *TUB3* or a control plasmid spotted onto rich media or rich media supplemented with 10 µg/ml benomyl. Plates were incubated at 30°C for 3 days before imaging. C. 10-fold dilution series of screen hits with the indicated *TUB1* allele spotted onto rich media or rich media containing 5 µg/ml benomyl. Plates were incubated at 30°C for 3 days or at 15°C for 17 days before imaging. D. Representative western blot of α-tubulin contribution assay. Blots were probed for α-tubulin and Zwf1 (G6PD) as a loading control. Bands corresponding to Tub1-6xHis and Tub3 are labeled. E. Quantification of α-tubulin contribution assay for the fraction of α-tubulin that corresponds to Tub1-6xHis. Dots and triangles represent different biological replicates and are colored based on experiment. Bars represent mean ± 95% CI. P-values are from a t test comparing wild type and mutant, specific p-values are listed in the text.

To confirm that these genes specifically promote the function of the *TUB1* intron, we used epistasis experiments to test whether the sensitivity of the null mutants to microtubule stress would be rescued by the *tub1Δ∷TUB3* allele, but not *tub1Δ∷TUB3^TUB1i^*. We find that the benomyl sensitivity of *ncl1*Δ mutants was not rescued by either allele, in fact, it is exacerbated (Figure 4C). In contrast, the benomyl sensitivity of both *swc3*Δ and *vps71*Δ are rescued by *tub1Δ::TUB3*, but not by *tub1Δ∷TUB3^TUB1i^*(Figure 4C). We find similar results for growth at low temperature (15°C), which represents a different microtubule-destabilizing stress (Figure 4C). These results are consistent with *SWC3* and *VPS71* operating in a pathway with the *TUB1* intron, and suggest that *NCL1* operates in a separate pathway.

Finally, we asked whether these genes and the intron selectively promote the expression of the Tub1 protein. Delineating the relative protein levels of the Tub1 and Tub3 isotypes on a denaturing gel is a challenge, since Tub1 contains 447 amino acids and Tub3 contains 445 amino acids. To improve resolution, we inserted a 6xHis tag between codons 43 and 44 of *TUB1*, allowing us to clearly separate the two isotypes on a 10% acrylamide gel (Figure 4D). Lysate from *tub3*Δ cells exhibits a single, slower migrating band, confirming that the slower-migrating band corresponds to Tub1-6xHis (Figure 4D). This method shows that in wild-type cells, 67% of α-tubulin is Tub1, which is consistent with previous data (Figure 4D, E; Bode et al., 2003; Gartz Hanson et al., 2016). When the *TUB1* intron is replaced with the *ACT1* intron, the amount of Tub1-6xHis is reduced to 50% of total α-tubulin (p = 0.0002, Figure 4D, E). Removal of the *TUB1* intron had a weaker effect on the fraction of α-tubulin that is Tub1-6xHis (p = 0.079, Figure 4D, E). This result suggests that the intron promotes the expression of Tub1 protein, and may play a role in regulating the balance between the two α-tubulin isotypes.

With this method we next asked if the genes identified in our screen maintain the ratio of Tub1-6xHis to Tub3 protein. *ncl1Δ* mutant cells show a small but significant reduction in the proportion of α-tubulin that is Tub1-6xHis, from 67% to 61% (p = 0.044, Figure 4B, C). Neither *swc3Δ* nor *vps71Δ* mutants show a difference in the ratio of the α-tubulin isotypes, compared to wild type (p = 0.79 and p = 0.22, Figure 4B, C). A null mutant in *CDC40*, which is known to disrupt the splicing of the *TUB1* intron, also failed to show a significant change in the ratio of the α-tubulin isotypes. As expected, *gim5Δ* mutant cells show a decrease in total α-tubulin, but no significant decrease in the ratio of the α-tubulin isotypes (p = 0.114; Figure 4D, E). Together these results suggest that some of the genes we identified through our screen may promote the expression of Tub1-6xHis and not Tub3, while other genes may be less important for isotype balance.

## Discussion

α-tubulin genes commonly contain 5’ introns, but how these introns impact expression and ensure sufficient α-tubulin production has not been established. In this study, we find that the intron within the budding yeast α-tubulin *TUB1* promotes α-tubulin expression and is important during microtubule stress. While we find that cells can survive without the intron, disrupting the intron sequence diminishes α-tubulin protein levels and response to microtubule stress. Finally, we performed an unbiased screen for genes that could promote α- tubulin expression through the *TUB1* intron. Our screen results also suggest a key role for α-tubulin introns in maintaining the balance between heterodimer production and turnover (Figure 5).

**Figure 5.**
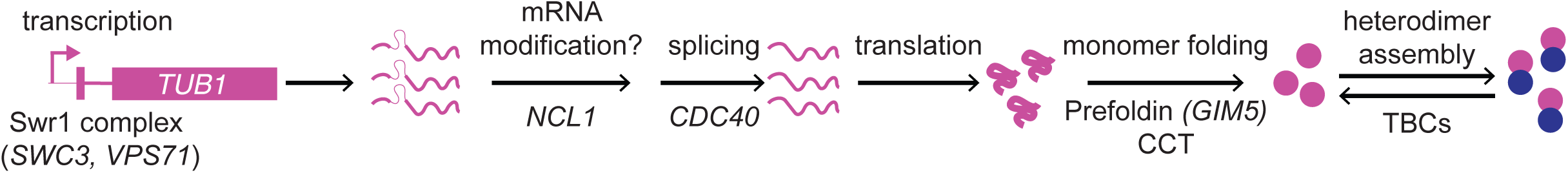
Proposed model for how the *TUB1* intron regulates α-tubulin expression. Diagram of α-tubulin biogenesis. Includes the genes we identified as acting through the *TUB1* intron for promoting α-tubulin expression and where we would expect they act in the pathway.

The *TUB1* intron appears to behave differently from the *ACT1* intron. The *ACT1* intron is known to promote transcription (Agarwal & Ansari, 2016; Furger et al., 2002; Moabbi et al., 2012), and simply inserting this intron into the coding sequence of another gene has been shown to boost gene expression in several cases (Agarwal & Ansari, 2016). However, the *ACT1* intron does not rescue the *TUB1* intron function in our experiments, and instead decreases the expression of *TUB1* (Figure 1C-E; 4D and E). We propose that the *TUB1* intron works together in a locus-specific manner that depends on either the promoter and/or the coding sequence. Consistent with this hypothesis, we find that combining the *TUB1* intron with the *TUB3* coding sequence is more resistant to benomyl than full replacement by *TUB3*, but less resistant than wild-type *TUB1*. These results suggest synergy between the *TUB1* promoter and the *TUB1* intron to promote α-tubulin expression, perhaps by the intron recruiting chromatin regulators to enhance transcription activation, which has been observed for introns in other *S. cerevisiae* genes (Moabbi et al., 2012).

Our genetic screen may provide clues to distinguish between these proposed functions and reveal key regulators. In our screen, we identified putative regulators of *TUB1* expression through the intron, using the criteria of hypersensitivity to microtubule stress (benomyl) and to loss of the alternative α-tubulin isotype (*tub3*Δ), and positive genetic interaction when combined with the *TUB1* allele that lacks the intron (*tub1Δi*). Our list of genes encompasses both genes that show no additive sensitivity to microtubule stress and genes where benomyl sensitivity is rescued when combined with *tub1Δi*. While there are various, previously reported functions among this set of 34 genes, we did find several RNA binding proteins, components of chromatin remodeling complex, and known regulators of tubulin biogenesis and turnover (Figure 3). *SWC3*, *VPS71* and other members of the Swr1 complex are likely to promote *TUB1* transcription either basally or in response to microtubule stress. Our data do not distinguish between these possibilities but do demonstrate clear specificity for the *TUB1* intron, consistent with the intron acting synergistically with the *TUB1* promoter (Figure 4, 5).

*CDC40*, a spicing factor, is known to be required for efficient splicing of *TUB1* pre-mRNA into mature mRNA (Figure 5; Burns et al., 1999, 2002). We propose that *NCL1*, a tRNA methyltransferase, may impact the translation of nascent α-tubulin through either post-transcriptional modifications of tRNAs or through unidentified modification of the *TUB1* mRNA itself (Figure 5). Together, this network of genes acts with the *TUB1* intron to promote α-tubulin function, which may be particularly important during microtubule stress.

Our screen also identified genes where ablating gene function appears to suppress the benomyl sensitivity caused by loss of the *TUB1* intron (25/33 genes; Table S2). Within this set are genes involved in tubulin biogenesis and turnover (Figure 3B). Specifically, we identified three members of the Gim/prefoldin complex that help fold nascent α- and β-tubulin monomers (*GIM4*, *PAC10*, *YKE2*), and three tubulin-binding cofactors (TBCs) that assemble and turn over α/β-tubulin heterodimers (*CIN1*, *CIN4*, *PAC2*; Gao et al., 1993, 1994; Kortazar et al., 2007; Tian et al., 1996). In particular, the human homologs of Cin1 and Pac2 are important for driving heterodimer turnover (Tian et al., 1999). How might loss of tubulin biogenesis and turnover ameliorate the effect of diminished α-tubulin expression by *tub1Δi*? We speculate that this may be explained slowing the production of folded β-tubulin and/or delaying the turnover of a diminished α/β- heterodimer pool. Our previous work established that yeast cells must maintain higher levels of α-tubulin compared to β-tubulin to prevent toxic accumulation of the latter (Wethekam & Moore, 2023). If diminished α- tubulin expression in *tub1Δi* cells creates an imbalance between α- and β-tubulin, then loss of β-tubulin folding by the Gim/prefoldin complex could help restore proper balance by decreasing β-tubulin levels. In addition, if diminished α-tubulin expression by *tub1Δi* creates a smaller pool of tubulin heterodimers, then turning down the rate of heterodimer turnover could increase the tubulin pool by extending the lifetime of tubulin proteins (Figure 5). Our data point towards a model where tubulin production and heterodimer turnover must be properly balanced to meet the cell’s demand for tubulin (Figure 5). This model will require further testing, including a better understanding of the mechanism(s) of tubulin turnover.

Finally, our study demonstrates different levels of activity for the 5’ introns in *TUB1* and *TUB3*, indicating that 5’ introns could be a point of regulation for creating blends of α-tubulin isotypes. We find that the *tub1*Δ*∷TUB3^TUB1i^* allele that replaces the *TUB3* intron with the intron from *TUB1* improves resistance to microtubule stress (Figure 2). This result suggests that while differences in amino acid sequence between Tub1 and Tub3 proteins may contribute to benomyl sensitivity, differences in intron sequence also contribute. In contrast, when we replace the *TUB1* intron with the *TUB3* intron we see a mild benomyl sensitivity suggesting that the *TUB3* intron does not fully replace the *TUB1* intron (Figure 2C). This level of phenotype is not different from that observed for cells lacking the *TUB1* intron (Figure 2C). Comparing the two intron sequences highlights several differences that could determine activity (Figure 2A). The first difference is that the *TUB3* intron is almost 3x as long as the *TUB1* intron (300bp for *TUB3* vs 118bp for *TUB1*) (Figure 2A, Schatz et al., 1986). Longer introns have been correlated with increased mRNA and protein expression; for example, the *ACT1* intron is 310bp and *ACT1* is highly expressed across the cell cycle (Blank et al., 2020; Juneau et al., 2006). Despite its longer intron, we find that there is less Tub3 protein in cells compared to Tub1 protein (Figure 4F; Bode et al., 2003; Gartz Hanson et al., 2016; Juneau et al., 2006). While intron length may influence function, it is not sufficient to determine expression levels in this case. The second key difference in the intron architecture is the branch point to 3’ splice site length (Figure 2A). For both *TUB1* and *ACT1*, the branch point to the 3’ splice site is 41 and 42 bp, respectively, and is consistent with the majority of introns in other yeast genes (Figure 2A; Cellini et al., 1986). *TUB3* on the other hand has a branch point to 3’ splice site length of 139 bp (Figure 2A). Longer branch point to 3’ splice site lengths increase the likelihood for alternative 3’ splice sites and require secondary structures to ensure appropriate 3’ splice site selection and efficient splicing (Cellini et al., 1986; Gahura et al., 2011). Within the *TUB3* intron, we identified several possible alternative 3’ splice sites, including one that is closer to the branch point (Figure S1). Splicing at these sites would lead to frameshifts and disrupt Tub3 translation; there is some evidence that an alternative 3’ splice site is used in *TUB3* (Kawashima et al., 2014). The relevance of these sites or other potential sites in regulating the production of Tub3 or total α-tubulin is still unknown. Our findings establish 5’ introns as important regulators α-tubulin gene function and a better understanding of their molecular functions may shed new light on conserved mechanisms that balance tubulin isotype expression.

## Methods

### Yeast Manipulation and Culturing

Yeast manipulation, media, and transformations were performed by standard methods (Amberg et al., 2000). Deletion mutants were generated by PCR-based, homologous recombination methods (Petracek & Longtine, 2002). All gene and intron swap alleles were generated by PCR amplification from plasmid templates described below (Table S3), and transformation and homologous recombination into diploid strains where one copy of *TUB1* is replaced by a hygromycin B resistance marker (yJM0591; Table S2; Goldstein & McCusker, 1999). Heterozygous diploids with one wild-type and one mutant allele were then dissected to acquire haploids. All alleles were confirmed by sequencing. To build the *tub1Δi* allele, the coding sequence from KGY2914 (Burns et al., 2002), was amplified by PCR and transformed into diploid strain yJM0591.

### Plasmid Construction

To build the plasmid containing a 6xHis tag inserted between codons 43 and 44 in *TUB1*, genomic DNA from strain yJM1796 was used as a template for a PCR amplicon containing 450 bp of *TUB1* UAS, the coding sequence and intron, and 450 bp of UTR including a *URA3* marker 280bp 3’ of the stop codon. This amplicon was cloned into pRS314 at the NotI and KpnI sites to create pJM738. To build the *tub1Δi* plasmid QuikChange Mutagenesis oligos were generated to remove the intron from pJM738 plasmid. To build the *tub1i∷ACT1i* and the *tub1i∷TUB3i* plasmids, a Gibson assembly reaction was used to combine pJM738 with the *ACT1i* or *TUB3i* amplified from the genomic DNA from yJM1837. To build the *tub1∷TUB3* locus swap plasmid, a Gibson assembly reaction was used to replace the *TUB1* coding sequence and intron in pJM738 with the *TUB3* coding sequence and intron, which was amplified from pJM886. To build the *tub1∷TUB3^TUB1i^* plasmid, first a Gibson assembly reaction was used to exchange the *TUB1* intron into the tub3 locus in plasmid pJM886. Then a second Gibson assembly reaction was used to replace the *TUB1* coding sequence and intron in pJM887 with the *tub1∷TUB3^TUB1i^*sequence.

### Doubling Time Measurement

Cells were grown in 3 mL of rich liquid media (YPD) to saturation at 30°C and diluted 50-fold into fresh media. The diluted cultures were then aliquoted into a 96-well plate, with three to six technical replicates per experiment, and incubated at 30°C while single orbital shaking in a plate reader. We used two different instruments for our experiments. A Cytation3 plate reader (BioTek, Winooski, VT) was used for experiments with strains: yJM1837, 1838, 0120, 0121, 4478, 4479, 4611, 4745 and Epoch2 microplate reader (BioTek, Winooski, VT) was used for experiments with strains: yJM1837, 0120, 5124-5127, 5158-5163. The OD_600_ was measured every 5 minutes for 24 hours. Doubling time was calculated by fitting the growth curves to a nonlinear exponential growth curve as previously published (Fees & Moore, 2018). Each experiment was repeated three independent times with wild-type cells included in each experiment as an internal control. P-values are from student’s t-test.

### Benomyl Sensitivity Assay

Cells were grown in rich liquid media to saturation at 30°C, and a 10-fold dilution series of each culture was spotted to either rich media plates or rich media plates supplemented with 5 or 10 µg/mL benomyl (Sigma Aldrich #381586, St. Louis, MO). Plates were grown at the indicated temperature for the indicated days.

### Western Blotting

Soluble protein lysates were prepared under denaturing conditions using the method of Zhang et al (Zhang et al., 2011). To make lysate log-phase cells were pelleted and resuspended in 2 M Lithium acetate and incubated for 5 minutes at room temperature. Cells were pelleted again and resuspended in 0.4 M NaOH for 5 minutes on ice. Cells were pelleted and resuspended in 2.5x Laemmli buffer and boiled for 5 minutes. Before loading gels, samples were boiled and centrifuged at 6,000xg for 3 minutes. Total protein concentration of clarified lysate was determined by Pierce 660 nm protein assay with the Ionic Detergent Compatibility Reagent before blotting (Cat. 1861426 and 22663, Rockford, IL). 0.75ug of total protein was loaded per lane for western blots looking to separate the two α-tubulin isotypes. Samples were run on 10% Bis-Tris PAGE gels in NuPAGE MOPS running buffer (50 mM MOPS, 50 mM TrisBase, 0.1% SDS, 1 mM EDTA, pH 7.7) at 0.04 mAmp per gel for 2.25 −2.5 hours to separate α-tubulin isotypes or 1.25 hours to determine the number of molecules per cell. Gels were transferred to PVDF (Millipore, IPFL85R) in NuPAGE transfer buffer (25 mM Bicine, 25 mM Bis-Tris, 1 mM EDTA, pH7.2) at 0.33 mAmp for 1 hour. Membranes were then blocked for 1 hour at room temperature in PBS blocking buffer (LI-COR, 927-70001). Membranes were probed in PBS blocking buffer including the following primary antibodies: mouse-anti-α-tubulin (4A1; at 1:100; Piperno and Fuller, 1985), mouse-anti-β- tubulin (E7; at 1:100; Developmental Studies Hybridoma Bank, University of Iowa), rabbi-anti-Zwf1 (Glucose-6- phosphate dehydrogenase; Sigma A9521; at 1:10,000) overnight at 4°C. After incubation in primary antibody, membranes were washed once in PBS for 5 minutes at room temperature and then probed with the following secondary antibodies: goat-anti-mouse-680 (LI-COR 926-68070, Superior, NE; at 1:15,000) and goat-anti-rabbit-800 (LI-COR 926-32211; at 1:15,000) for 1 hour at room temperature. After incubation in secondary antibodies, blots were washed twice in PBST (1XPBS, 0.1% Tween-20), once in PBS, and imaged on an Odyssey Imager (LI-COR, 2471).

### Quantifying Tubulin Concentration

To determine levels of tubulin in the cell, wild-type or mutant cells were grown to log phase in rich media at 30°C. To prepare lysate of 5×10^7^ cells in 50 µL samples, cells from each culture were counted on a hemocytometer and the appropriate volume of cells was determined based on the hemocytometer counts and prepared as described above. To confirm the number of cells per volume of culture, separate samples of cells from the overnight cultures were diluted and plated to rich media at approximately 200 cells/plate. After 2 days at 30°C, the number of colonies/plate was counted, and the fraction of cells that formed colonies was determined. Lysate was resuspended in 2.5x Laemmli buffer and standards of purified yeast tubulin (Wethekam and Moore, 2023) were prepared by diluting protein to 2.5 ng/µL in 2.5x Laemmli buffer. Samples containing increasing amounts of cells (3.5, 4.5, 6, and 8 x 10^6^) or purified tubulin (4,10, 15, 30 and 40 ng of total protein heterodimers or 2, 5, 7.5, 15, and 20 ng of α-tubulin) were loaded and blotted as described above. Band intensities were quantified using the gel analysis plug-in in FIJI.

To confirm the amount of cell lysate loaded per lane, we used a method described in Wethekam & Moore (Wethekam & Moore, 2023). Zwf1 loading control intensity was plotted against expected number of cells loaded and the r^2^ value was calculated. If the r^2^ value was <0.75, the outlier lane was identified, and we excluded it. If at least three lanes from a replicate did not generate an r^2^ value ≥ 0.8, then that replicate was removed. The proportionality of Zwf1 signal to cell number was determined by dividing the measured Zwf1 signal intensity per lane by the number of expected cells loaded, the average of those values represents the Zwf1 signal per cell for that blot. That value was used to recalculate the number of cells in each lane by dividing the Zwf1 band intensity by the average Zwf1 signal per cell.

Finally, we converted band intensities for α-tubulin from cell lysates into estimated nanograms of protein using a standard curve ranging from 2.0 to 20 ng of α-tubulin. Linearity of the signal was assessed as a function of ng loaded for each standard curve and set a cut off of r^2^ < 0.85. In some cases we found that higher amounts (i.e. 20 ng) of pure tubulin deviated from the linear regression and in these cases we limited the standard curves to lower amounts of tubulin. We used these linear regressions to calculate the ng of α-tubulin in each lane of cell lysate for that blot. The calculated ng of α-tubulin was then divided by the estimated number of cells in that lane, and converted to molecules/cell with the following:

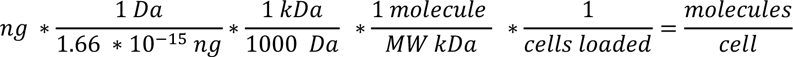

The molecule/cell values for biological replicates were averaged for a single experiment and the corresponding α-tubulin were compared to determine the ratio of α-tubulin. P-values are from student’s t-test after a one-way ANOVA with a Tukey post-hoc test for p < 0.05.

### Deletion Collection and Synthetic Genetic Array Screen

The yeast deletion collection was prepared as described in Baryshinkova et al (2010) (Baryshnikova et al., 2010). Briefly, the deletion collection (Open Biosciences now Horizon Discovery, YSC1053, Gardner & Jaspersen, 2014; Winzeler et al., 1999) was thawed from −80°C storage and transferred onto rich media + 50 µg/mL G418 with the Singer RoToR (Singer Instruments, Somerset, UK). Four, 96-well plates were condensed onto one plate resulting in 384 colonies and then amplified to quadruplicate to generate an array of 1536 deletion mutants per plate. These plates were incubated at 30°C, and then transferred to four different plates with the following conditions: rich media incubated at 30°C, rich media incubated at 15°C, rich media with 2% DMSO incubated at 30°C, and rich media with 10 µg/ml benomyl and 2% DMSO incubated at 30°C. Plates were incubated at 30°C for two days, or at 15°C for ten days. Each plate was scanned on an Epson Perfection V300 Photo scanner and processed using the gitter plugin in R as described on http://gitter.ccbr.utoronto.ca (Wagih & Parts, 2014). Plates were manually rotated in FIJI to ensure colony recognition. Once colonies were identified, data were normalized and scores were calculated using http://sgatools.ccbr.utoronto.ca (Wagih et al., 2013). Since this analysis compares the deletion collection on different conditions, no linkage was taken into account for scoring. Each of the conditions was normalized to the rich media alone, but we also compared the 10 µg/mL benomyl + 2% DMSO data to the 2% DMSO data to identify genes that are specifically sensitive to benomyl and not DMSO. We used a score > −0.08 and p-value < 0.05 as the cutoffs for genes we considered benomyl sensitive. The scores from that comparison were used for analysis in Figure 3.

To screen for genetic interactions with the deletion collection, we generated a *tub1Δi* allele in strain Y7092 (Baryshnikova et al., 2010). We carried out the Synthetic Genetic Array screen following the protocol described in Baryshinkova et al. (2010) (Baryshnikova et al., 2010). 1536-colony array plates containing haploid, double mutants of *tub1Δi* and the deletion mutants were stamped to four different conditions: rich media incubated at 30°C, rich media to be incubated at 15°C, rich media with 2% DMSO incubated at 30°C, and rich media with 10 µg/ml benomyl and 2% DMSO incubated at 30°C. Plates were incubated at 30°C for two days, or at 15°C for ten days. All of the plates were processed as described for the deletion collection alone with the following modifications: i) for each condition, double mutants were compared to the deletion collection alone; ii) we completed this analysis for both 200 kb of linkage and no linkage (Baryshnikova et al., 2010; Wagih et al., 2013).

### GO Term Analysis and Plotting

To obtain enriched GO Terms for both hits identified in the genetic screen and the negative genetic interactors pulled from the Saccharomyces Genome Database we used g:Profiler (Raudvere et al., 2019). All settings were left at default with the exceptions of: species switched to *S. cerevisiae* and all results were displayed. For Figure 3A, the −log_10_(p-values) were plotted for GO Terms with p <0.05 that were also non-redundant with other GO Terms.

### Protein Interaction Networks

Network map showing previously annotated protein-protein interactions were generated using GeneMania (Montojo et al., 2010). Network branches are weighted by Cellular Component GO terms.

### α-tubulin Isotype Analysis by Western Blot

To measure the proportional levels of each α-tubulin isotype we used strains where a 6X histidine tag was inserted between codons 43 and 44 of *TUB1*, allowing the resolution of Tub1 and Tub3 proteins by western blot. Western blots were then analyzed using the following method. We first rotated the gel image in FIJI to ensure that each lane would be square to our ROIs. To sample the band intensity in each lane, we used the Gel Analysis Plugin to create four 5-pixel by 500-pixel ROIs and place them along the vertical axis of the gel and across the middle of each lane, approximately 20, 40, 60, and 80% of the width of the lane. Intensity profiles were plotted and the background for each ROI was removed by drawing a line at the minimum intensity value. To identify the peak for Tub1 and Tub3 lines were drawn in 3 places: on either side of the α-tubulin signal and then at the minimum intensity point between the Tub1 and Tub3 peaks. We then measured the area within each peak corresponding to the slower-migrating Tub1 with the internal 6X histidine tag and faster migrating Tub3. The fraction of Tub1 reported in Figure 4C represents the intensity value of Tub1 divided by the sum of Tub1 and Tub3 from the same sample. Each dot in Figure 4E represents a biological replicate as the mean from the 4 positions in a single lane.

## Acknowledgments

We are grateful to current and former members of the Moore lab for advice and discussions on this project. We would like to thank Blossom Lee for her assistance with the genetic screen, Dr. Kathy Gould for sharing the *tub1Δi* strain, Dr. Michael McMurray for lending his Singer RoToR for the genetic screen, and Dr. Jay Hesselberth and Saylor Strugar for guidance with RNA biology. This work was supported by NIH R35 GM 136253 to J.K.M. L.C.W. was supported by T32 GM136444 and the Bolie Scholar Award. The authors declare no competing financial interests.

---

**Figure S1.**
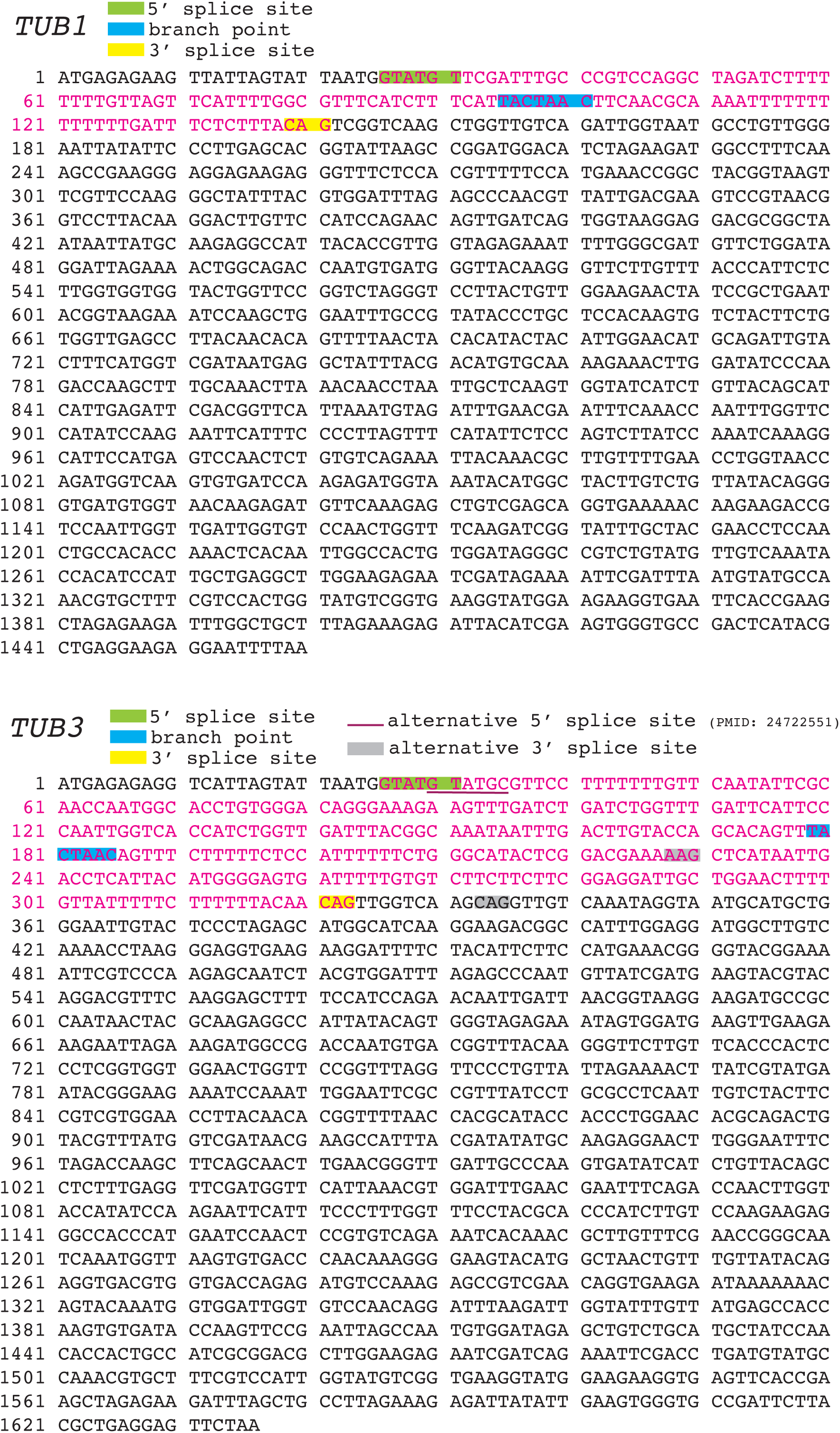
Annotated coding and intron sequences for *TUB1* and *TUB3*. Sequences retrieved from the Saccharomyces Genome Database showing the coding sequence (black) and intron sequence (pink). 5’ splice sites are highlighted in green, branch points in blue, and 3’ splice sites in yellow. *TUB3* includes a previously identified alternative 5’ splice site and two possible alternative 3’ splice sites (Kawashima et al., 2014).

**Table S1. Benomyl Sensitive Genes**

Table of genes with a score > −0.08 and p < 0.05 on 10 µg/mL benomyl + 2% DMSO compared to 2% DMSO alone. Contains scores, standard deviation, and p-value for each gene.

**Table S2. Genetic screen results**

Table of our genetic screen results. Table includes growth scores, standard deviation and p-value on 10 µg/mL benomyl + 2% DMSO compared to 2% DMSO alone, interaction with TUB3 and associated PMID, scores, standard deviation, and p-value for deletion with tub1Δi compared to deletion alone on 10 µg/mL benomyl + 2% DMSO. Final column describes if the score ± the standard deviation overlaps between the deletion on benomyl alone and the deletion with tub1Δi.

**Table S3.**
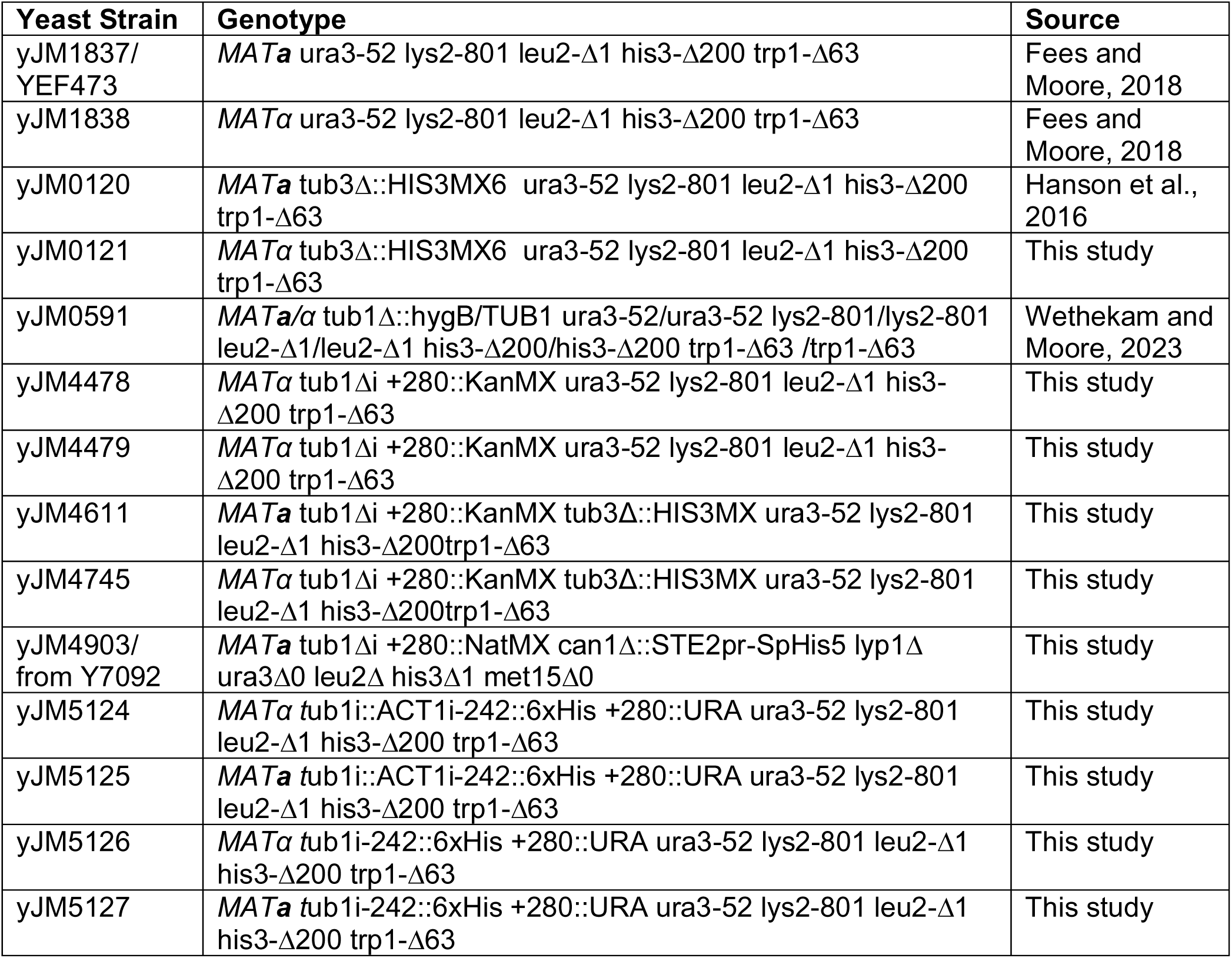
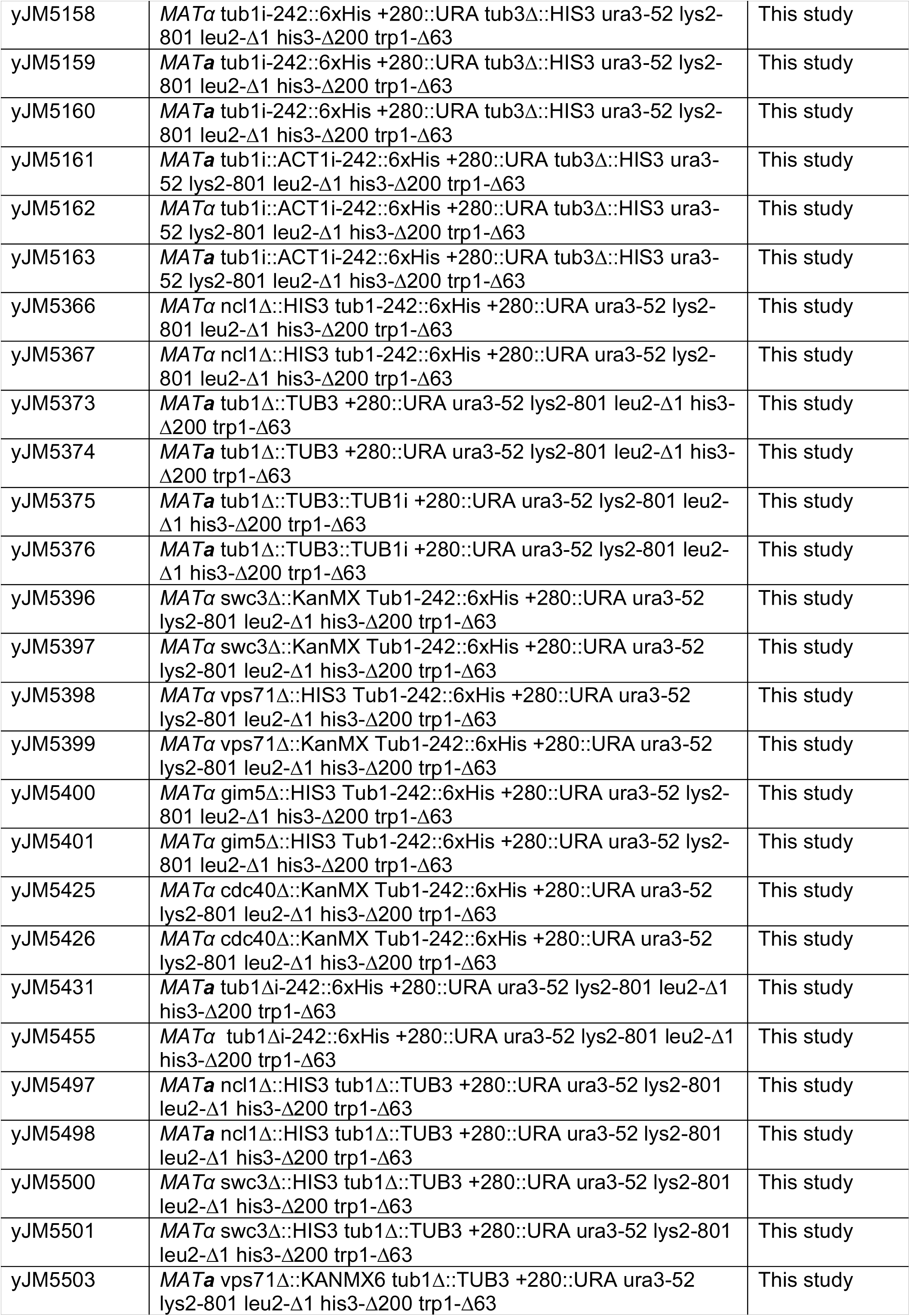
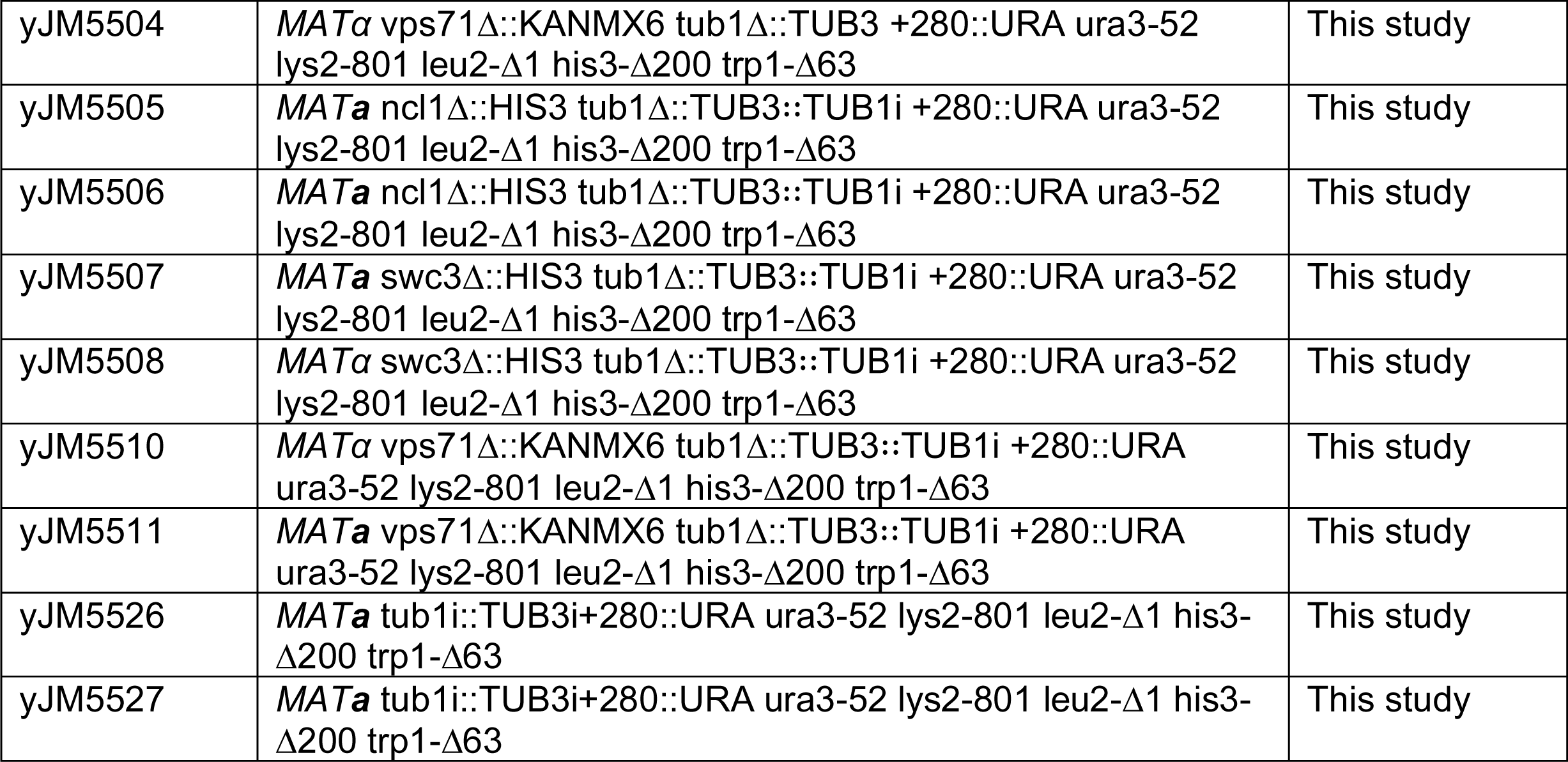
Yeast Strains.

**Table S4.** Plasmids.

| Plasmid ID | Plasmid Name | Source |
| --- | --- | --- |
| pJM0738 | UAS-TUB1 +243::6xHis +280::URA in pRS314 | This study |
| pJM0822 | UAS-tub1i::ACT1i -242::6xHis-UTR +280::URA in pRS314 | This study |
| pJM0823 | UAS-tub1i::TUB3i -242::6xHis-UTR +280::URA in pRS314 | This study |
| pJM0886 | UAS-TUB3-UTR in pRS316 | This study |
| pJM0887 | UAS-tub3i::TUB1i-UTR in pRS316 | This study |
| pJM0927 | UAS-tub1::TUB3-UTR +280::URA in pRS314 | This study |
| pJM0928 | UAS-tub1::TUB3<sup>TUB1i</sup>-UTR +280::URA in pRS314 | This study |
| pJM0949 | UAS-tub1Δi-UTR +280::URA in pRS314 | This study |

